# Parental death destabilizes wolf packs despite the benefits of larger pack size

**DOI:** 10.64898/2026.10.15.759480

**Authors:** C. Aaron Bott, Roblyn Brown, Daniel R. MacNulty

**Author notes:** Corresponding Author: *C. Aaron Bott*, Department of Wildland Resources and Ecology Center, Utah State University, Logan, Utah 84322, USA.

## Abstract

1. Cooperative breeding is a fundamental feature of many animal family groups. However, few studies have quantified whether variation in group size meaningfully buffers social and reproductive consequences following parental death.
2. Using 17 years of data from a gray wolf (*Canis lupus*) population in Oregon, USA, we evaluated how parental death (the death of one or both parents) influenced pack dissolution, reproductive failure, and pup recruitment, and whether larger pack sizes mitigated these effects.
3. Across 303 pack-years, we recorded 69 parental deaths, 39 pack dissolutions, and 112 reproductive failures. Packs that experienced parental death were nearly five times more likely to dissolve (95% CI: 2.12–11.12), nearly 16 times more likely to fail reproductively (95% CI: 5.34– 47.26), and recruited 66% fewer pups (95% CI: 49–78% fewer) than packs in which both parents survived.
4. Larger packs were associated with improved demographic outcomes overall, but these group-size benefits were insufficient to fully offset the elevated risks associated with parental death.
5. Our results show that numerous individuals can provide generalized pack-level demographic resilience but do not functionally replace parental roles, highlighting the importance of evaluating social structure in addition to population size.
6. By clarifying the limits of social buffering in a cooperatively breeding carnivore, this study advances understanding of how social systems respond to parental death.

## Introduction

Animal sociality shapes patterns of survival, reproduction, and evolutionary fitness (Wilson, 1975; Hamilton, 1964; Silk, 2007). Singular cooperative breeding is a social system in which a dominant parental pair monopolizes reproduction while non-breeding auxiliary members (“helpers”) assist in raising offspring (Emlen, 1995; Lukas & Clutton-Brock, 2012; Smith et al., 2017). This system, observed in birds, mammals, and fish, relies on biparental care, kin-based cooperation, and social cohesion to maintain reproductive output and group stability (Cornwallis et al., 2010; Lukas & Clutton-Brock, 2018). Auxiliary members can enhance reproductive success, breeder fitness, and group persistence (Ausband & Mitchell, 2021; Morrison et al., 2021), but such benefits vary with population density, group maturity, and the availability of replacements (Clutton-Brock, 2002; García-Ruiz et al., 2022; Ausband & Bassing, 2024).

In cooperative breeders, the demographic consequences of group size are often framed in terms of group augmentation, where individuals in larger groups experience higher survival or reproductive success, and Allee effects, where very small groups suffer reduced fitness or increased extinction risk (Kingma et al., 2014; Rubenstein & Abbot, 2017; Thorley et al., 2023). These patterns suggest that larger groups may be better able to absorb the loss of important group members because more auxiliary individuals remain to contribute to offspring care and other cooperative functions. However, parental care can make substantial contributions to offspring survival and reproductive performance (Pilakouta et al., 2018), raising the alternative possibility that the specialized functions of parents cannot readily be replaced by additional group members. Thus, whether larger group size buffers the demographic consequences of parental death may depend on the extent to which auxiliary members can functionally compensate for the roles performed by parents. For species of conservation concern, where populations may be socially fragile, distinguishing between these alternatives is particularly important (Markham et al., 2015; Lukas & Clutton-Brock, 2018; Maldonado-Chaparro & Chaverri, 2021). Hereafter, we use “parent” synonymously with breeder, as identified in previous studies (e.g., Mech, 1999).

Here, we use a 17-year dataset on a gray wolf (*Canis lupus*) population in Oregon, USA, to evaluate these alternative hypotheses. Wolves are well-suited for this test because they live in family groups (“packs”) anchored by a parental pair (Mech, 1999) that usually mates for life (Milleret, 2017; Mech, 2024). Under a social-buffering hypothesis, larger packs should be better able to compensate for parental death because more auxiliary members remain to contribute to cooperative functions, consistent with the generalized demographic benefits of larger group size. Alternatively, under a parental-irreplaceability hypothesis, larger packs may perform better overall but fail to specifically buffer parental death because parents perform specialized reproductive and social functions that other pack members cannot readily assume. Wolf parents jointly provision young, defend territories, and coordinate group movement (Mech & Boitani, 2003; Mech, 2000; Stahler et al., 2013). Long-term parental pair bonds may also improve reproductive performance through accumulated experience, coordination, and familiarity with territories and resources (Ausband, 2019). Because parents and auxiliary members can occupy distinct reproductive and social roles in cooperative groups, remaining pack members may not be functionally equivalent to parents (Lukas & Clutton-Brock, 2018; Ausband & Bassing, 2024). Previous studies have shown that parental loss can reduce reproduction and pack persistence in wolves (Brainerd et al., 2008; Borg et al., 2015; Cassidy et al., 2023), but whether larger group size mitigates these consequences remains unclear.

We quantified the effects of parental death on three key processes—pack dissolution, reproductive failure, and pup recruitment. We predicted that parental death would increase pack dissolution and reproductive failure and reduce pup recruitment. The two alternative hypotheses generated contrasting predictions about the role of pack size. Under the social-buffering hypothesis, the demographic consequences of parental death should be offset by the generalized benefits associated with larger pack size. Under the parental-irreplaceability hypothesis, larger pack size should improve demographic performance overall, but these benefits should be insufficient to fully compensate for parental death. We therefore evaluated the effects of parental death and pack size on each demographic outcome and assessed whether the benefits associated with larger pack size were sufficient to offset the consequences of parental death.

Finally, because wolf recovery and management are frequently evaluated using population-level metrics, we conducted a secondary analysis to evaluate whether these pack-level effects were detectable at the broader population scale. We hypothesized that the demographic consequences of parental death would be attenuated at the population level because statewide population growth integrates the dynamics of many packs, allowing losses in some packs to be offset by reproduction, recruitment, and persistence elsewhere in the statewide population. Accordingly, we predicted that annual parental deaths would have a weaker association with statewide population growth than the strong associations predicted for demographic outcomes within individual packs.

## Materials and methods

Oregon’s diverse landscapes—including high deserts, temperate conifer forests, and mountains— support varied ecological conditions that influence wolf dispersal, survival, and reproduction. Wolves recolonized Oregon naturally in the early 2000s following extirpation, aided by reintroductions in the Northern Rocky Mountains during 1995–1996. Oregon’s population began recovering *via* immigration rather than translocation (Oregon Department of Fish and Wildlife (ODFW), 2019).

Following partial delisting from the federal Endangered Species Act in 2011, wolf management became dual-jurisdictional: state-managed in eastern Oregon and federally protected in western Oregon (Figure 1). Recreational hunting and trapping are prohibited statewide; human-caused mortality primarily reflects illegal killing (poaching) or targeted agency control actions implemented in response to chronic livestock depredation.

**Figure 1.**
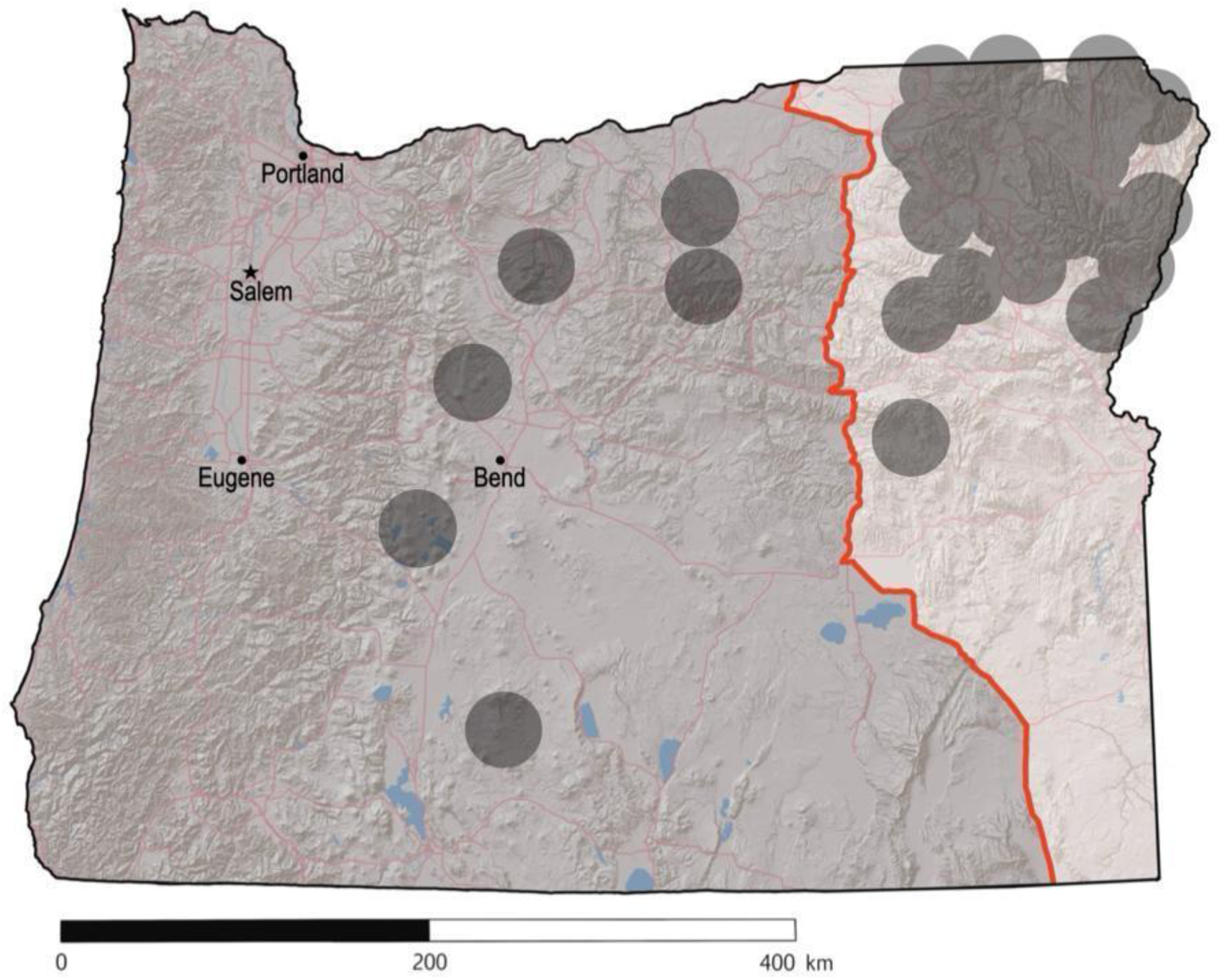
The study area encompassed the entire state of Oregon, USA. Since 2011, wolves located in the unshaded area to the right of the red line (Highway 395/78/95) are managed independently by the Oregon Department of Fish and Wildlife, whereas wolves in the shaded region to the left of the red line are federally listed under the Endangered Species Act. Recreational hunting and trapping of wolves are prohibited throughout the state. Dark dots indicate parental death locations. Base map: Esri World Shaded Relief (© Esri, USGS, NOAA); map modified by authors.

**Figure 2.**
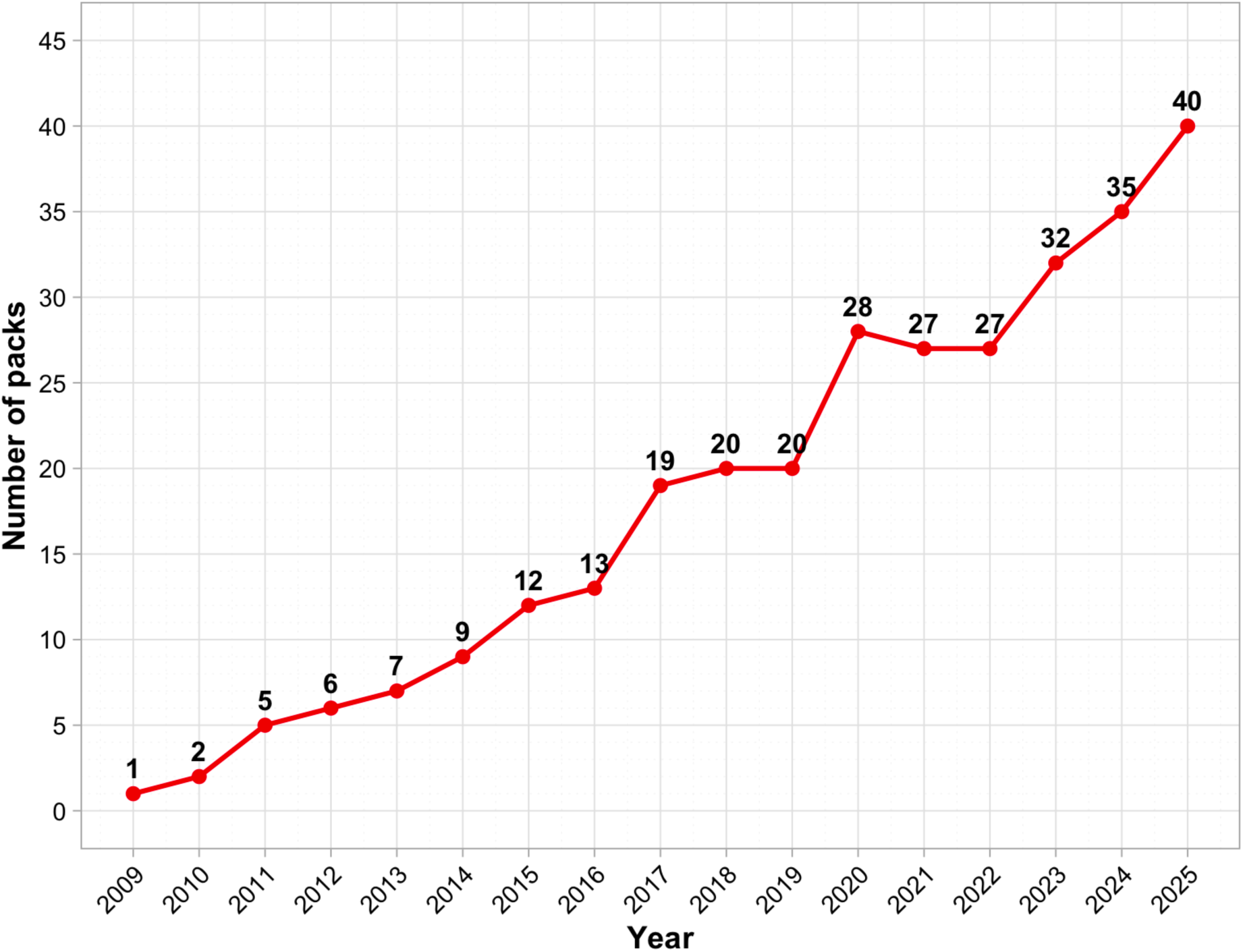
Annual number of wolf packs in Oregon, USA, during the study period, 2009-2025.

### Data collection and monitoring

We monitored wolves from 2009 to 2025 using radio-collaring and non-invasive methods to identify parents and quantify pack composition and size. Captured wolves were fitted with very high frequency (VHF) or global positioning system (GPS) collars following aerial darting in winter or foothold trapping during spring–fall. The monitoring program sought to maintain at least one radio-collared wolf in each pack whenever possible (ODFW, 2019). During the study, 215 wolves were radio-collared and >90% of packs were radio-collared at some point. Remote cameras were also deployed throughout known wolf territories each summer. Intensive monitoring from January through mid-March included repeated camera surveys, ground and aerial telemetry, and direct observations, providing repeated detections of packs and individuals and high-confidence assessments of pack persistence, composition, and reproduction. Animal capture and handling were conducted as part of ODFW wildlife management activities and did not require Institutional Animal Care and Use Committee (IACUC) approval or separate state research permits.

Telemetry, cameras, and genetic sampling provided complementary information. GPS and VHF telemetry tracked packs and individuals through time, whereas cameras documented pack composition, reproductive status, and social associations. Approximately 450 genetic samples were obtained from captured (n = 215) and dead (n = 235) wolves, with additional opportunistic samples from scat, and were used to confirm individual identity and parentage when available.

We identified parent wolves using physical traits (e.g., nipple development, testis size; Mech, 2006; Barber-Meyer & Mech, 2015), genetic information, and repeated evidence of denning, provisioning, or association with pups. Individuals were distinguished by radio-collar status, pelage and other physical characteristics, and repeated observations through time. Long-term pack associations further aided consistent identification of parents (Mech, 1999). These methods were implemented within ODFW’s broader statewide wolf monitoring program (ODFW, 2019). Pups were identified using cameras, telemetry-assisted observations, and aerial surveys based on body size, physical characteristics, dentition when handled, behavior, and association with parents. The biological year began each April to align with denning and pup production.

We defined a pack as a social group of ≥2 wolves that included a confirmed breeding pair with documented reproductive history at the start of the biological year. This ensured that packs had demonstrated reproductive potential before any parental death. Pairs observed during winter surveys (January–mid-March) generally represented remnants of previously larger packs rather than newly formed pairs and were retained to distinguish social dissolution from simple numerical decline. A pack was considered persistent if ≥2 socially affiliated wolves remained together at year’s end, as indicated by repeated observations, telemetry, or cameras; packs reduced to a single wolf or no longer detected within their known territory or range were classified as dissolved.

Deaths were confirmed through field investigations, radio-collar mortality signals, necropsies, and carcass recovery through agency investigations, public reports, or routine monitoring. Causes were classified as human-caused, natural, or unknown; human-caused mortality included illegal killing and targeted agency control. Parental deaths associated with full-pack agency removals were excluded to avoid conflating social collapse with intentional population reduction. Whenever possible, parental death and pack composition were corroborated using multiple sources, including telemetry, carcass recovery, genetics, cameras, and repeated field observations.

Wolf packs generally center on a single breeding pair, although plural breeding occasionally occurs (Ausband, 2018; Mech, 2024). We documented <5 packs with more than one reproducing female during the study, precluding robust analysis of plural breeding strategies. Standardized a priori criteria were applied to pack persistence, reproductive failure, and dissolution. Territories reoccupied ≥1 year after dissolution were classified as new packs when social continuity could not be established, reducing the risk of pseudo-replication.

Our objective was to evaluate pack-level demographic responses associated with parental death rather than broader ecological drivers of wolf demography. Pack size, measured before each reproductive season, represented the immediate social environment and a biologically meaningful proxy for group augmentation. Population density increased monotonically during the study and covaried strongly with year and recovery phase; these effects were partially accommodated by including year as a random effect. Analyses therefore focused on demographic outcomes associated with parental death irrespective of cause.

### Predictor variables

The primary predictor was parental death, coded as binary (0 = no parental death, 1 = ≥1 parental death). As sensitivity analyses, we evaluated two alternative mortality metrics. First, documented non-parental death was coded as binary (0 = none, 1 = ≥1) to assess whether demographic responses were specific to parental rather than general pack-member mortality. Because non-parental deaths are more difficult to document consistently, this variable included only deaths confirmed through radio-collars, agency lethal action, field investigation, or carcass recovery. Second, parental death was classified as none, single-parent, or biparental to evaluate responses to severity of parental loss. We explored effects of parent sex and mortality timing, but sample sizes within individual categories were insufficient for reliable estimation, and these variables were excluded from final models.

Pack size was measured at the start of the biological year as the number of wolves counted before the reproductive season; lone wolves and transient non-breeders were excluded. Although detection error is possible, most packs included at least one radio-collared wolf and were repeatedly surveyed using multiple methods, making systematic bias in pack-size estimates unlikely.

### Statistical analysis Pack dissolution

We used a logistic generalized linear mixed model (GLMM) with a binomial distribution and logit link to evaluate the effect of parental death on pack dissolution. Dissolution was defined as the disappearance of a pack or its reduction to a single individual by the end of the following biological year. Dissolution was assessed over a one-year interval, from the end of one annual survey to the next (January–mid-March), consistent with our biological calendar. Parental death was the primary predictor, with pack size as a covariate. Random intercepts for pack and year accounted for repeated measures and annual variation. We also evaluated a first-order autoregressive [AR(1)] correlation structure within packs across years. Although the model converged, parameter estimates and variance components were unstable, and inference differed substantially from the simpler random-intercept model. Because diagnostic evaluation suggested overparameterization and poor reliability, the simpler model was retained. Odds ratios (ORs) and 95% confidence intervals (CIs) were calculated. Model diagnostics using DHARMa residual simulations indicated appropriate model fit with no evidence of zero-inflation.

### Reproductive failure

Reproductive failure was modeled with a logistic GLMM including parental death and pack size as fixed effects and random intercepts for pack and year. Reproductive failure was defined as fewer than two pups surviving by the time of annual monitoring (January–mid-March), with pups typically born in mid-April. Analyses were restricted to packs that persisted through the entire biological year. We evaluated an AR(1) correlation structure within packs across years to account for temporal autocorrelation, but this structure did not materially improve model performance and was not retained. ORs and 95% CIs were reported.

### Pup recruitment

Pup counts were modeled using a negative binomial GLMM to account for overdispersion. Recruitment was measured as the minimum pup count observed during annual surveys (January–mid-March) in packs that remained intact through the full year. Parental death and pack size were fixed effects, with random intercepts for pack and year. We evaluated an AR(1) correlation structure within packs across years, but the model exhibited convergence problems and produced unstable variance estimates without materially altering fixed-effect inference. The simpler negative binomial GLMM was therefore retained. Zero-inflation was detected (DHARMa, p < 0.001), but the model was retained given our focus on inference for fixed effects. Incidence rate ratios (IRRs) were reported.

### Interaction effects

To evaluate whether the association between parental death and each demographic outcome varied with pack size, we additionally fit models that included an interaction between parental death and pack size. These models otherwise retained the same response distributions, fixed effects, and random-effects structures as the corresponding primary models. Because parental deaths were uncommon in larger packs, particularly those containing >8 wolves, interaction estimates at larger pack sizes were based on relatively sparse data. We therefore interpreted interaction estimates cautiously and retained the additive models for primary inference.

### Sensitivity analyses

To evaluate the robustness of our primary results, we conducted two sets of sensitivity analyses for each demographic response. First, we added documented non-parental deaths as an additional predictor to assess whether demographic effects were specific to parental death. Second, we replaced the binary parental death variable with a three-level factor distinguishing no parental death, single-parent death, and biparental death. These analyses were not intended to replace the primary models, but rather to evaluate whether the direction and magnitude of inferred parental death effects were sensitive to alternative mortality classifications.

### Population growth and parental death

Although not a primary objective, we explored whether parental death was associated with variation in statewide wolf population growth beyond density dependence. We fit Bayesian state-space models (SSM) to annual wolf population estimates in Oregon from 2009–2025, modeling population growth as a function of previous population size (log N) and parental death. To assess robustness to scaling and collinearity with population size, parental death was expressed using three formulations: total annual parental deaths (raw), per-capita parental deaths, and residualized parental deaths derived from a regression of deaths on log N. Models were fit both with and without density dependence to evaluate sensitivity to model structure. For all formulations, posterior summaries were used to estimate baseline population growth (b₀), density dependence (b₁), parental-death effects (b₂), process variance (σ_process), and observation variance (σ_observation).

We assigned normal priors to the initial latent population state [x₁ ∼ Normal(log(20), 1)] and regression coefficients (b₀, b₁, and b₂ ∼ Normal(0, 1)), and gamma priors to the process and observation precisions [τ_process and τ_observation ∼ Gamma(0.001, 0.001)]. Models were run using three MCMC chains of 60,000 iterations each, with the first 30,000 iterations discarded as burn-in and every 10th post-burn-in sample retained.

### Software

All analyses were conducted in R version 4.0.2 (R Core Team, 2020) using the *survival* (Therneau, 2020), glmmTMB (Brooks et al., 2017), DHARMa (Hartig, 2020), and ggeffects (Lüdecke, 2018) packages. SSMs were fit using the NIMBLE package (de Valpine et al., 2017) in R via MCMC sampling. Convergence and mixing were assessed using Gelman–Rubin diagnostics (Gelman & Rubin, 1992) and effective sample sizes. Across all SSM formulations and monitored parameters, the minimum effective sample size was 1,676 and the maximum Gelman–Rubin statistic (R^) was 1.01 (maximum upper confidence limit = 1.02), indicating satisfactory convergence and mixing.

## Results

We recorded 39 pack dissolutions and 112 reproductive failures during 303 pack-years from 2009 to 2025. A total of 69 parental deaths were documented in 54 events; 37 (68.5%) were due to human causes, 4 (7.4%) to natural causes, and 13 (24.1%) were classified as unknown. We observed 39 single-parent death events (22 female and 17 male) and 15 biparental death events. Wolf pack sizes ranged from 2 to 15 individuals, with a mean of 6 wolves per pack; 83% of packs contained eight or fewer wolves— within the range reported for other western U.S. wolf populations (Fuller et al., 2003; Smith et al., 2020). Only 17 of the 303 pack-years (5.6%) ended with exactly two wolves, the minimum threshold for our pack definition. Most of these two-wolf groups were verified remnants of larger packs earlier in the same biological year (see *Data collection and monitoring*). Three pack-years involving complete agency removal of all pack members were excluded from demographic analyses because pack disappearance in those cases reflected management action rather than natural social dynamics.

### Pack dissolution

Packs that experienced at least one parental death were nearly five times more likely to dissolve within a one-year interval than packs in which both parents survived (OR = 4.86, 95% CI: 2.12–11.12, p < 0.001; Table 1; Figure 3). Each additional pack member reduced the odds of dissolution by 21% (OR = 0.79, 95% CI: 0.66–0.95, p = 0.010), though no pack size guaranteed persistence. Model diagnostics indicated a good fit with no evidence of zero-inflation.

**Figure 3.**
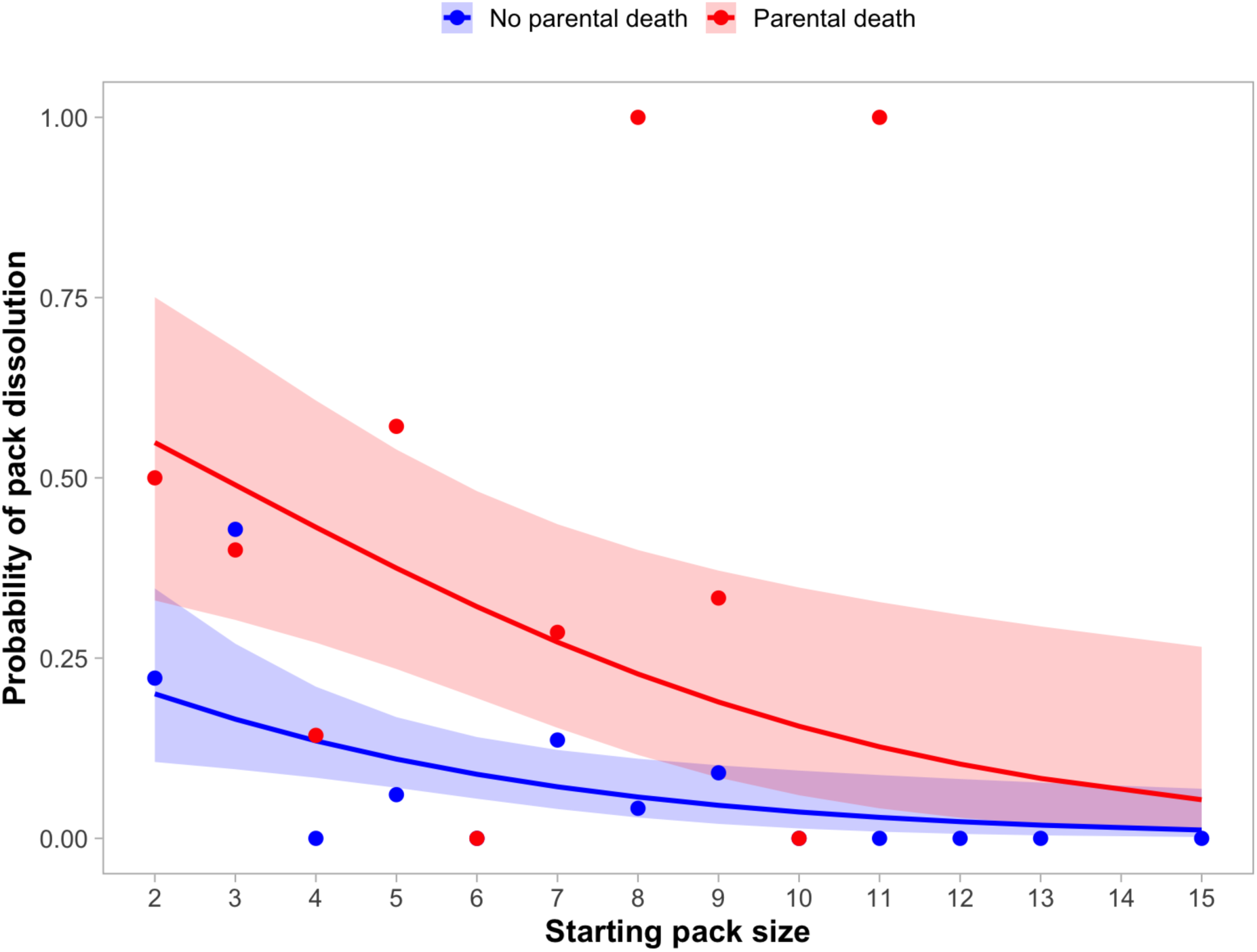
Effects of pack size and parental death (i.e., the death of one or both parents) on the probability of pack dissolution. Dissolution was defined as the loss of social persistence, such that fewer than two socially affiliated wolves remained together by the end of the biological year. Curves depict predicted probabilities from a logistic generalized linear mixed model, with shaded areas representing 95% confidence intervals. Red indicates packs that experienced parental death, whereas blue indicates packs in which both parents survived. Points represent observed proportions of pack-years that dissolved for each combination of starting pack size and parental-death status.

**Table 1.** Results from a logistic generalized linear mixed model assessing the effects of parental death (i.e., the death of one or both parents) and pack size on the probability of wolf pack dissolution in Oregon, USA. Estimate refers to the log-odds coefficient; Std. Error is its standard error; z is the Wald test statistic; and p is the associated p-value. Odds ratios (OR) indicate how the odds of dissolution change with each predictor: values above 1 reflect increased odds, values below 1 reflect reduced odds. The 95% confidence intervals (Lower–Upper 95% CI) show the likely range of values for each odds ratio.

| Predictor | Estimate | Std. Error | z | p | OR | Lower 95% CI | Upper 95% CI |
| --- | --- | --- | --- | --- | --- | --- | --- |
| (Intercept) | -0.91 | 0.52 | -1.70 | 0.089 | 0.40 | 0.14 | 1.15 |
| Parental death | 1.58 | 0.42 | 3.74 | <0.001 | 4.86 | 2.12 | 11.12 |
| Pack size | -0.24 | 0.09 | -2.56 | 0.010 | 0.79 | 0.66 | 0.95 |

### Reproductive failure

Packs that experienced at least one parental death were nearly 16 times more likely to experience reproductive failure than those where both parents survived (OR = 15.89, 95% CI: 5.34–47.26, p < 0.001; Table 2; Figure 4). Each additional pack member reduced the odds of reproductive failure by 15% (OR = 0.85, 95% CI: 0.73–0.98, p = 0.030). Model diagnostics indicated a good fit with no evidence of zero-inflation.

**Figure 4.**
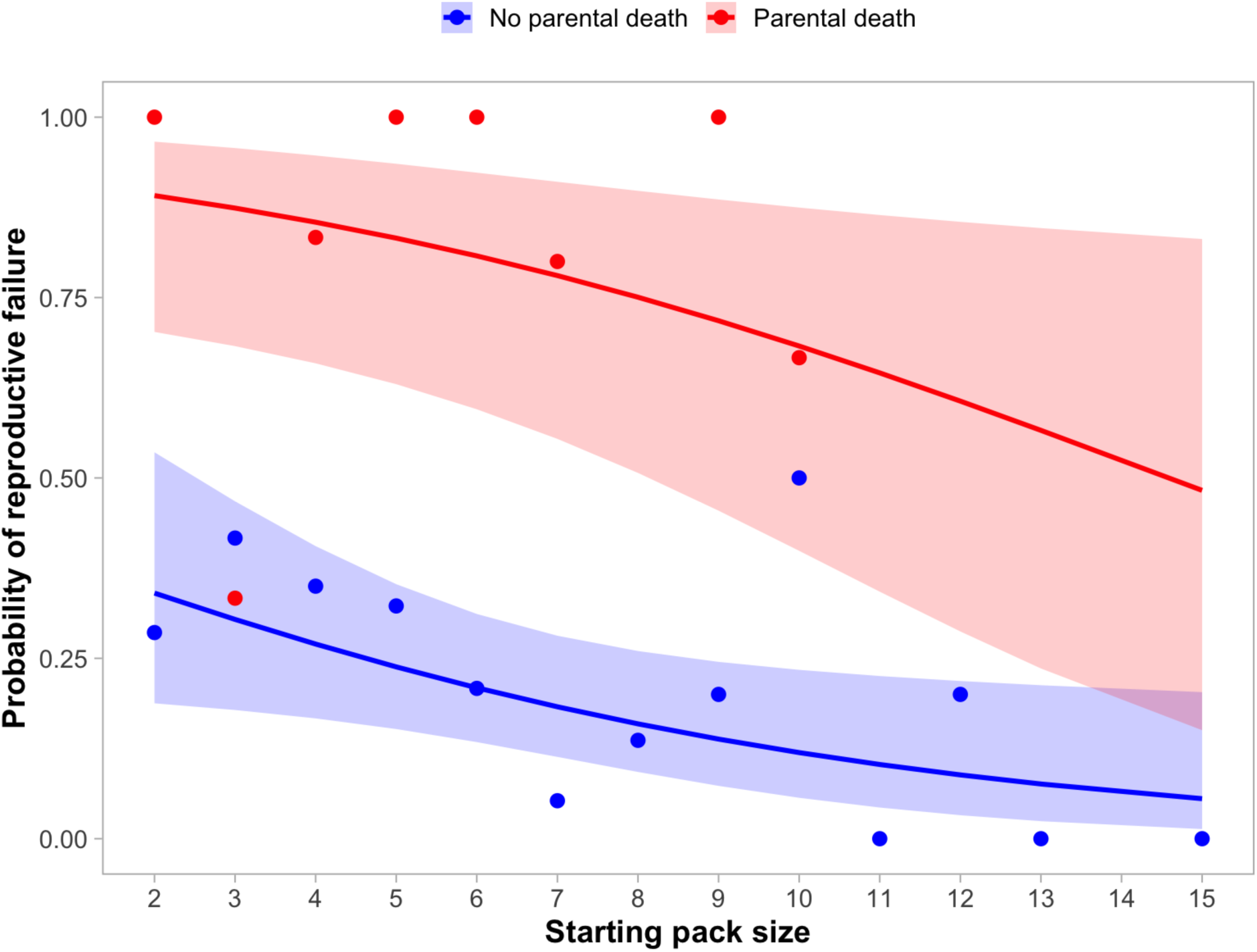
Effects of pack size and parental death (i.e., the death of one or both parents) on the probability of reproductive failure, defined as ≤1 pup surviving to the annual winter survey (January–mid-March). Curves depict predicted probabilities from the fitted model, with shaded areas representing 95% confidence intervals. Red indicates packs that experienced parental death, whereas blue indicates packs in which both parents survived. Points represent observed proportions of pack-years that experienced reproductive failure for each combination of starting pack size and parental-death status.

**Table 2.** Results from a logistic generalized linear mixed model assessing the effects of parental death (i.e., the death of one or both parents) and pack size on the likelihood of reproductive failure in wolf packs in Oregon, USA. Reproductive failure was defined as fewer than two pups surviving by the time of annual monitoring (January–mid-March), with pups typically born in mid-April. Estimate refers to the effect of each predictor on the log-odds scale; Std. Error is the standard error of the estimate; z is the Wald test statistic; and p is the associated p-value. OR (odds ratio) indicates how the odds of reproductive failure change with each predictor: values above 1 indicate increased odds, while values below 1 indicate decreased odds. The 95% confidence intervals (Lower and Upper 95% CI) show the likely range of values for each odds ratio.

| Predictor | Estimate | Std. Error | z | p | OR | Lower 95% CI | Upper 95% CI |
| --- | --- | --- | --- | --- | --- | --- | --- |
| (Intercept) | -0.33 | 0.53 | -0.61 | 0.541 | 0.72 | 0.25 | 2.06 |
| Parental death | 2.77 | 0.56 | 4.97 | <0.001 | 15.89 | 5.34 | 47.26 |
| Pack size | -0.17 | 0.08 | -2.17 | 0.030 | 0.85 | 0.73 | 0.98 |

### Pup recruitment

Among persisting packs, parental death reduced pup recruitment by 66% on average (IRR = 0.34, 95% CI: 0.22–0.51, p < 0.001; Table 3; Figure 5). Larger packs recruited more pups, but the effect was modest and not statistically significant (IRR = 1.04, 95% CI: 0.99–1.09, p = 0.11). Zero-inflation was detected, but alternative model structures produced convergence issues and did not materially change fixed-effect estimates; therefore, the negative binomial model was retained for inference.

**Figure 5.**
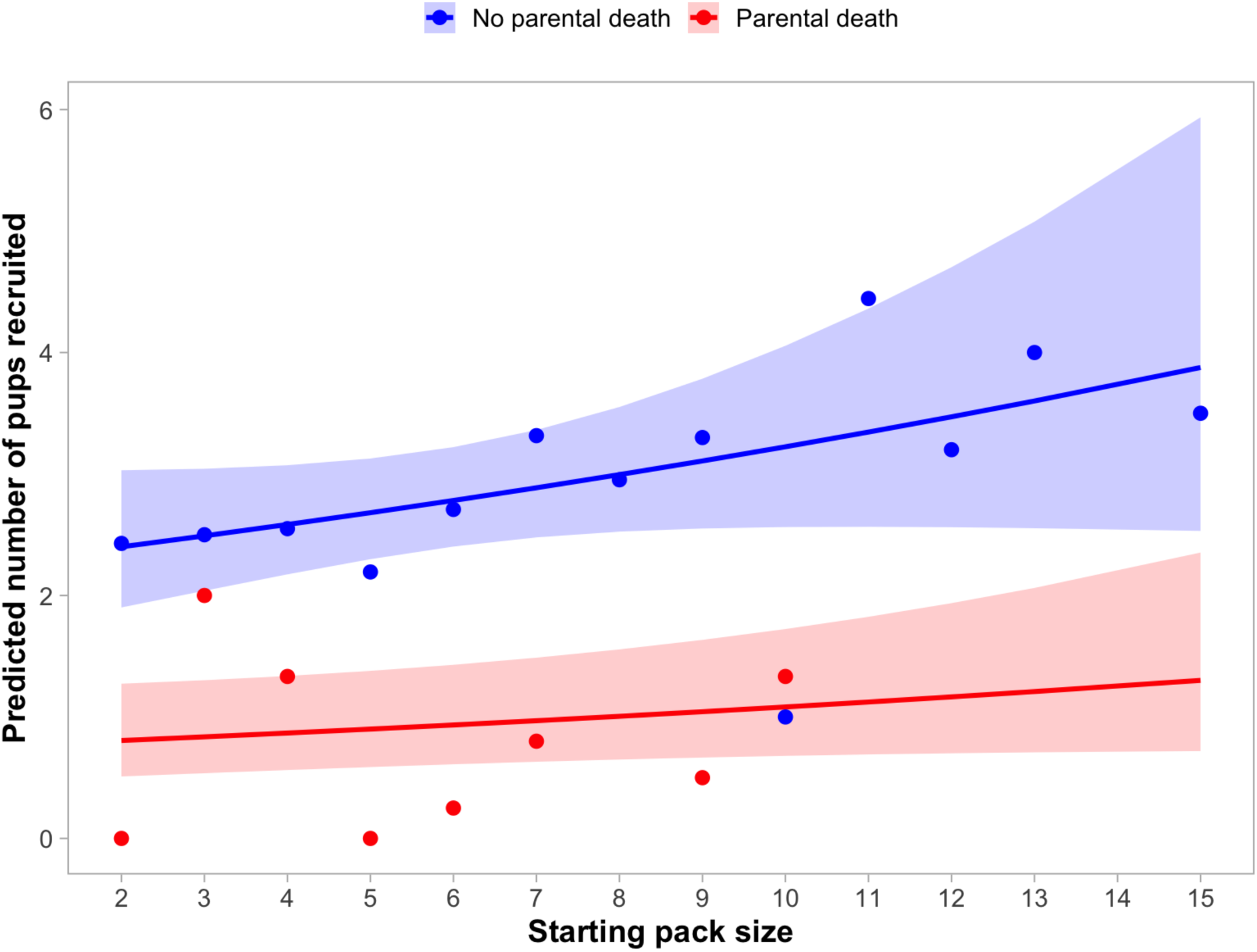
Effects of pack size and parental death (i.e., the death of one or both parents) on pup recruitment, measured as the minimum number of pups observed during annual winter surveys (January– mid-March). Curves depict model-predicted pup recruitment from a negative binomial generalized linear mixed model, with shaded areas representing 95% confidence intervals. Red indicates packs that experienced parental death, whereas blue indicates packs in which both parents survived. Points represent the observed mean number of recruited pups for each combination of starting pack size and parental-death status.

**Table 3.** Results from a negative binomial generalized linear mixed model evaluating the effects of parental death (i.e., the death of one or both parents) and pack size on pup recruitment in wolf packs in Oregon, USA. The response variable was the minimum number of pups surviving to 9-10 months. Estimate refers to the effect of each predictor on the log scale; Std. Error is the standard error of the estimate; z is the Wald test statistic; and p is the associated p-value. IRR (incidence rate ratio) indicates how the expected number of surviving pups changes with each predictor: values above 1 mean more pups are expected, while values below 1 mean fewer. The 95% confidence intervals (Lower and Upper 95% CI) show the likely range of values for each IRR.

| Predictor | Estimate | Std. Error | $z$ | $p$ | IRR | Lower 95% CI | Upper 95% CI |
| --- | --- | --- | --- | --- | --- | --- | --- |
| (Intercept) | 0.80 | 0.16 | 5.11 | <0.001 | 2.23 | 1.64 | 3.03 |
| Parental death | -1.09 | 0.21 | -5.11 | <0.001 | 0.34 | 0.22 | 0.51 |
| Pack size | 0.04 | 0.02 | 1.62 | 0.106 | 1.04 | 0.99 | 1.09 |

### Sensitivity analyses

Sensitivity analyses yielded results that were qualitatively similar to those of the primary models. Including documented non-parental death as a covariate did not materially alter the estimated effects of parental death on pack dissolution, reproductive failure, or pup recruitment (Tables S1, S3, and S5). Non-parental death was not associated with pack dissolution (OR = 1.03, 95% CI: 0.40–2.68, p = 0.94; Table S1) or pup recruitment (IRR = 0.83, 95% CI: 0.66–1.04, p = 0.10; Table S5), but was associated with increased odds of reproductive failure (OR = 2.43, 95% CI: 1.10–5.37, p = 0.029; Table S3). Analyses distinguishing single-parent and biparental mortality yielded effect estimates in the predicted direction, but biparental mortality estimates were highly uncertain because of limited sample sizes (Tables S2, S4, and S6).

### Interaction effects

Interaction models provided evidence that the association between pack size and dissolution differed depending on whether parental death occurred (β = 0.42, 95% CI: 0.04–0.80). The interaction model modestly improved fit relative to the additive dissolution model (ΔAIC = 2.74; likelihood-ratio test, p = 0.029). Predicted dissolution risk declined with increasing pack size when both parents survived but remained relatively constant following parental death, indicating that the protective association of larger pack size was attenuated following parental loss. However, parental-death events were sparse among larger packs, resulting in considerable uncertainty in estimates of the interaction at larger pack sizes. Interaction estimates for reproductive failure (β = 0.30, 95% CI: −0.19–0.79) and pup recruitment (β = −0.09, 95% CI: −0.28–0.10) were also uncertain. We therefore retained the additive models for primary inference and interpreted the interaction analyses cautiously.

### Population growth and parental death

Annual wolf population growth exhibited strong density-dependent regulation across all SSM formulations, with growth rates declining as population size increased (Tables 5 and 6). Posterior estimates for density dependence were consistently negative (b₁ = −0.16 to −0.17), and 95% credible intervals excluded zero in all full models.

**Table 4.**
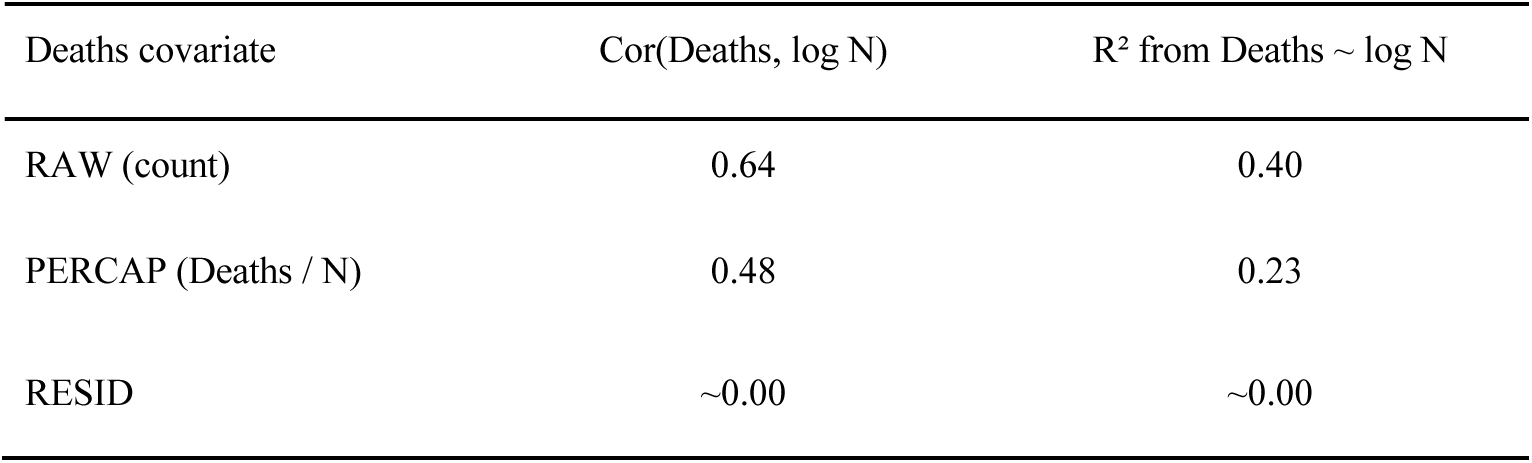
Collinearity between parental-death covariates and population density (log N) prior to standardization. Alternative covariate formulations were evaluated to assess the robustness of parental-death effects in population growth models.

| Deaths covariate | Cor(Deaths, log N) | R<sup>2</sup> from Deaths ~ log N |
| --- | --- | --- |
| RAW (count) | 0.64 | 0.40 |
| PERCAP (Deaths / N) | 0.48 | 0.23 |
| RESID | ~0.00 | ~0.00 |

**Table 5.**
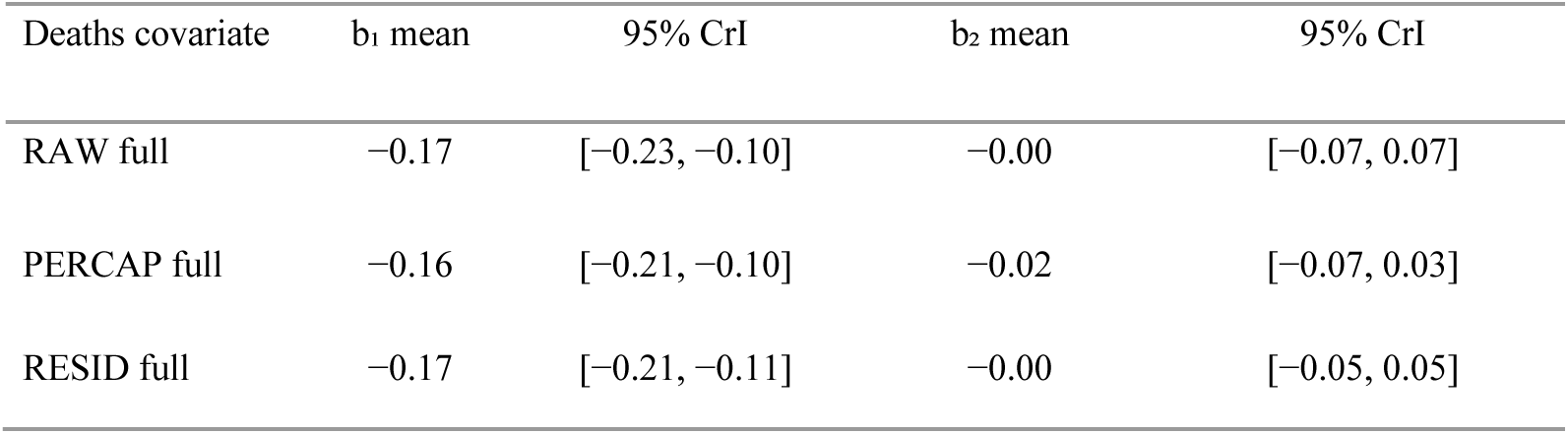
Posterior means and 95% credible intervals from Bayesian state-space models of Oregon wolf population growth including density dependence (b₁; log N) and parental-death effects (b₂), evaluated using alternative formulations of parental mortality.

**Table 6.** Posterior summaries from Bayesian state-space models of Oregon wolf population growth (2009–2025) incorporating density dependence (log N) and parental-death covariates. Values are posterior means (SD) with 95% credible intervals.

| Parameter | RAW deaths | PERCAP deaths | RESID deaths |
| --- | --- | --- | --- |
| $b_0$ (baseline growth) | 0.16 (0.02)<br>[0.13, 0.20] | 0.16 (0.02)<br>[0.13, 0.20] | 0.16 (0.02)<br>[0.13, 0.20] |
| $b_1$ (density dependence) | -0.17 (0.03)<br>[-0.23, -0.10] | -0.16 (0.03)<br>[-0.21, -0.10] | -0.17 (0.02)<br>[-0.21, -0.11] |
| $b_2$ (parental deaths) | 0.00 (0.03)<br>[-0.07, 0.07] | -0.02 (0.03)<br>[-0.07, 0.03] | 0.00 (0.03)<br>[-0.05, 0.05] |
| $\sigma_{\text{process}}$ | 0.07 (0.02)<br>[0.03, 0.12] | 0.06 (0.02)<br>[0.02, 0.11] | 0.07 (0.02)<br>[0.03, 0.12] |
| $\sigma_{\text{observation}}$ | 0.05 (0.02)<br>[0.02, 0.09] | 0.05 (0.02)<br>[0.02, 0.09] | 0.05 (0.02)<br>[0.02, 0.09] |

After accounting for density dependence and process variation, parental death did not explain additional variation in annual population growth. Estimated effects of parental deaths were consistently small and uncertain, with 95% credible intervals overlapping zero regardless of whether death was expressed as raw counts, per-capita rates, or residualized values (Tables 5 and 6; Figure 6). Posterior means for parental death effects ranged from −0.02 to 0.00, indicating little evidence that annual variation in parental death influenced statewide population growth once density dependence was considered.

**Figure 6.**
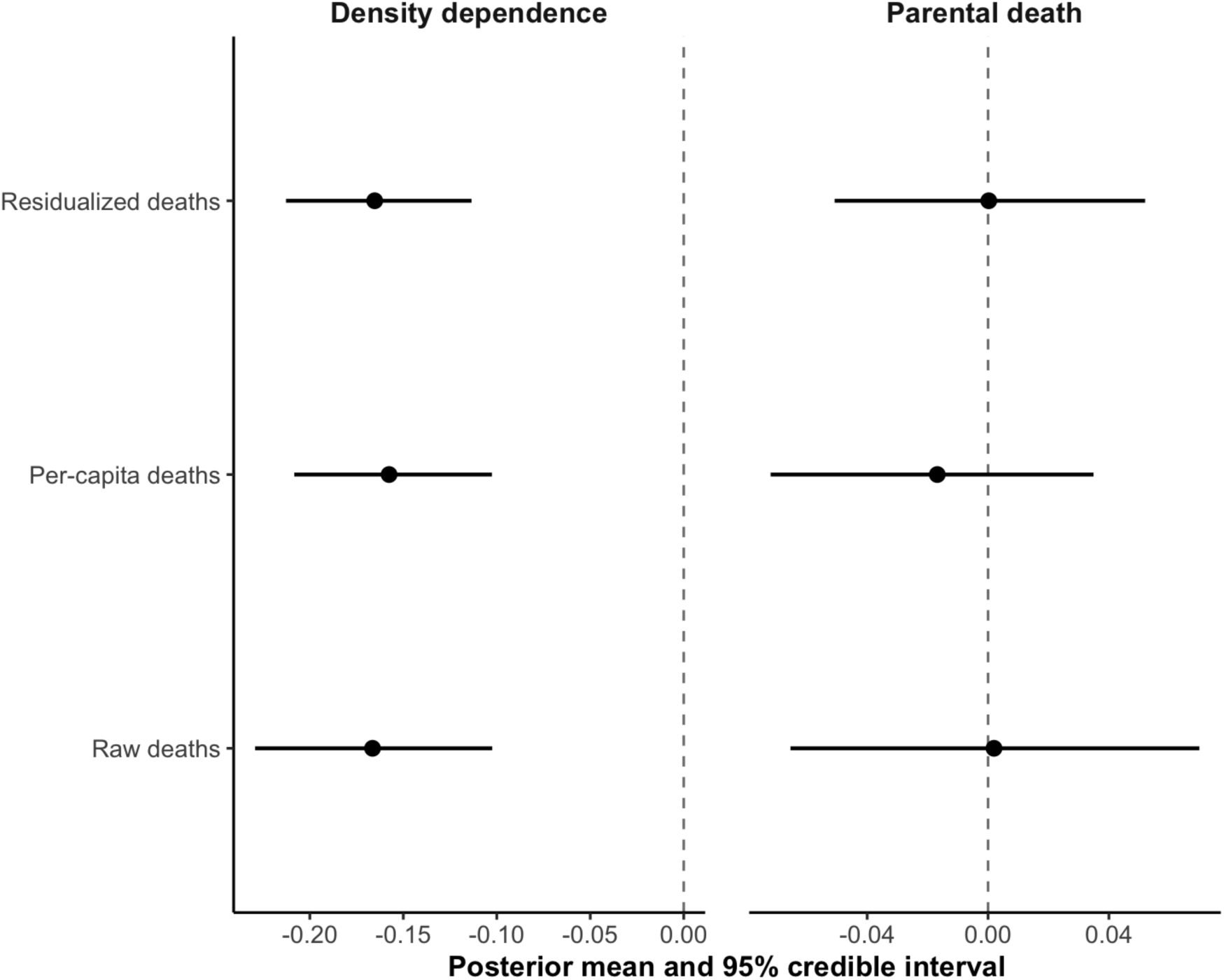
Posterior estimates from full state-space models of annual wolf population growth. Density dependence was consistently negative across model formulations, whereas parental death effects were small and uncertain, with 95% credible intervals overlapping zero. Error bars show 95% credible intervals.

Collinearity diagnostics showed that raw parental deaths covaried positively with population size (Table 4), whereas per-capita scaling reduced this association, and residualization effectively removed it. Consistent with these patterns, apparent negative associations between population growth and parental death in simplified models were attenuated once density dependence was included in the full models (Tables 5 and 6). The consistency of results across all three mortality formulations suggests that the absence of a detectable population-level effect was robust to alternative representations of parental death.

## Discussion

Parental death increased the odds of pack dissolution and reproductive failure and reduced pup recruitment, underscoring the importance of parental roles within wolf family groups (Emlen, 1995; Clutton-Brock, 2019). While larger pack sizes were associated with better pack-level outcomes across all measures, the benefits reflected generalized demographic resilience rather than functional replacement of parental roles. Each additional pack member reduced the odds of dissolution by 21%, reproductive failure by 15%, and was associated with a 4% increase in pup recruitment, although this last effect was not statistically significant (p = 0.11). During our study, pack sizes ranged from 2 to 15 individuals, with a mean of 6 wolves per pack; 83% of packs contained eight or fewer wolves. Model estimates indicated that the level of pack enlargement required to fully offset elevated dissolution or reproductive failure risk following parental death exceeded the size of most observed packs and, for pup recruitment, would require group sizes larger than those observed in our system. Thus, while larger groups provided modest demographic benefits consistent with group augmentation, they were generally insufficient to fully compensate for the demographic consequences of parental death. These results indicate that parental roles in wolves are not readily replaced by additional group members, even in relatively large social groups. Sensitivity analyses indicated that documented non-parental mortality showed a weaker and less consistent relationship with demographic outcomes than parental mortality. Although non-parental mortality was associated with reproductive failure, it was not associated with pack dissolution or pup recruitment (Tables S1, S3, and S5).

Our results were more consistent with the parental-irreplaceability hypothesis than with the social-buffering hypothesis. Although larger packs exhibited greater demographic resilience overall, we found little evidence that increasing pack size compensated for the demographic consequences of parental death. For dissolution specifically, the protective association of larger pack size was attenuated following parental death, although interaction estimates became increasingly uncertain at larger pack sizes because data for parental deaths were uncommon in these groups. The benefits provided by auxiliary pack members therefore appear not to be functionally equivalent to the roles performed by parents. This result is consistent with the specialized reproductive and social roles occupied by parents within wolf packs. Long-term parental pair bonds may enhance reproductive performance through accumulated experience, coordination, and familiarity with territories and resources (Ausband, 2019), while breeders and auxiliary members can occupy distinct reproductive and social roles in cooperative groups (Lukas & Clutton-Brock, 2018; Ausband & Bassing, 2024). More broadly, the consequences of losing socially important individuals vary widely among group-living species because social roles and cooperative functions are distributed differently among group members. Across birds and mammals, alloparental care varies substantially in both form and extent, while experimental and observational studies show that removal or loss of group members can reduce cooperative care, destabilize social relationships, fragment groups, or, in some systems, strengthen relationships among those remaining (Ben Mocha et al., 2023; Maldonado-Chaparro & Chaverri, 2021). For example, breeder turnover in dwarf mongooses (*Helogale parvula*) can trigger further social instability and reduced reproductive success (Kern et al., 2023), whereas mountain gorillas (*Gorilla beringei beringei*) provide a striking contrast: relationships with other group members strengthen following maternal loss and may buffer otherwise expected fitness costs (Morrison et al., 2021). Taken together, these contrasting responses suggest that resilience to the loss of important individuals depends less on group size alone than on the degree of functional redundancy within the social system. In wolves, additional pack members appear to provide generalized demographic benefits without fully replacing the specialized reproductive and social functions of parents. Taken together, our results are consistent with the concept of parents functioning as keystone individuals whose influence on group dynamics is disproportionate to their numerical contribution (Modlmeier et al., 2014), highlighting wolves as a useful system for examining the limits of social buffering in cooperatively breeding groups with singular reproduction.

To evaluate whether strong pack-level effects scaled up to detectable changes in statewide population dynamics, we conducted exploratory SSM analyses of annual wolf population growth. Oregon’s wolf population exhibited clear density-dependent regulation across all SSM formulations (Tables 5 and 6). After accounting for density dependence and process variation, estimated parental-death effects on annual population growth were small and uncertain across raw, per-capita, and residualized formulations (Tables 5 and 6). In models that excluded density dependence, raw parental deaths showed a negative association with population growth, but this effect was attenuated once density dependence was included, consistent with the covariance between parental deaths and population size in a growing population (Table 4). Overall, these exploratory analyses indicate that annual statewide growth is strongly influenced by density feedback and stochasticity, while any independent association with parental death is difficult to identify from this short time series despite strong pack-level demographic consequences.

This apparent decoupling of pack-level disruption from population-level growth suggests that continued numerical expansion can occur despite repeated parental deaths because population growth reflects the combined demographic performance of many packs rather than the stability of any individual family group. Consequently, reproductive losses or pack dissolution in some packs may be offset by successful reproduction, recruitment, and pack persistence elsewhere in the population (Creel & Rotella, 2010; Borg et al., 2015; Schmidt et al., 2017).

Importantly, the absence of a detectable population-level signal does not imply that parental death is biologically inconsequential to cooperative breeders. Rather, it highlights the scale-dependent nature of demographic processes in socially structured species: parental death was consistently associated with family group destabilization and reduced reproductive performance locally (Brainerd et al., 2008; Ausband, Mitchell, Stansbury, et al., 2017; Cassidy et al., 2023), but these effects may be masked at broader spatial scales where population growth reflects the aggregate dynamics of many packs at different stages of establishment and turnover (Fuller et al., 2003; Borg et al., 2015; Smith et al., 2020). Population-level growth metrics alone may therefore fail to capture biologically meaningful variation in social organization, even when numerical recovery criteria are met (Wallach et al., 2009; Maldonado-Chaparro & Chaverri, 2021; vonHoldt et al., 2025).

These findings matter for two reasons. First, they contribute to a broader understanding of how cooperative societies respond to the loss of reproductively and socially important individuals. Cooperative breeding is characterized by assistance from non-breeding group members, but the presence of helpers does not necessarily imply functional redundancy among group members. In cooperatively breeding mammals, reproduction is typically strongly skewed toward a dominant pair while non-breeding helpers contribute to offspring care (Lukas & Clutton-Brock, 2012). Our results demonstrate that these functions can remain demographically distinct: auxiliary members and larger group sizes enhanced general demographic resilience, but did not compensate for the loss of parental function. This distinction between helping and replacing may therefore be important for understanding the limits of social buffering in cooperative societies. Second, because wolves are managed using both numerical and social criteria, our findings have direct relevance for applied population assessment. Legal recovery thresholds frequently require not only a minimum abundance of wolves but also a minimum number of breeding pairs or packs (ODFW, 2019; USFWS, 1987). Consequently, wolf populations can meet numerical targets while simultaneously experiencing erosion of social units, particularly if parents are repeatedly killed (Wallach et al., 2009; vonHoldt et al., 2025), an insight relevant to management decisions involving targeted removals for conflict reduction (Brainerd et al., 2008; Borg et al., 2015; Cassidy et al., 2023).

Small sample sizes limited our ability to evaluate sex-specific effects or variation associated with mortality timing. Because our analyses focused on annual demographic outcomes, deaths during denning, pup-rearing, or other critical periods may have had different effects than those occurring later in the year. In addition, our analyses were designed to evaluate pack-level demographic responses associated with parental death rather than to model the full suite of ecological and management factors that may influence pack performance. Thus, although parental death was consistently associated with pack dissolution, reproductive failure, and reduced pup recruitment, we cannot fully distinguish the effects of parental death itself from correlated environmental or management conditions that may also affect demographic outcomes. Moreover, our study system represents a population without recreational hunting and trapping; responses in harvested populations may differ. Nonetheless, evidence from multiple regions indicates that parental turnover destabilizes packs across a range of ecological and management contexts (Brainerd et al., 2008; Borg et al., 2015; Ausband, Mitchell, Stansbury, et al., 2017; Cassidy et al., 2023; Ausband & Bassing, 2024), suggesting that our findings capture a general feature of wolf social organization rather than an Oregon-specific anomaly. Future studies integrating genetic pedigrees, behavioral data, and cross-population comparisons will be critical for determining how variation in social structure mediates resilience across cooperative breeding systems (Ausband, Mitchell, Stansbury, et al., 2017; Ausband, Mitchell & Waits, 2017).

More broadly, the capacity for groups to buffer the loss of parents or other socially important individuals should vary across cooperative breeding systems because the roles performed by group members differ among species and social systems. Across birds and mammals, cooperative breeders vary widely in reproductive skew, the prevalence and type of alloparental care, group composition, and the ecological conditions under which helpers remain with breeders (Thorley et al., 2023; Ben Mocha et al., 2023). Likewise, responses to social disruption range from fragmentation and reduced cooperation to compensatory strengthening of relationships among surviving group members (Maldonado-Chaparro & Chaverri, 2021; Morrison et al., 2021). These differences suggest that group size alone is an incomplete measure of social resilience: additional members should provide greater buffering when cooperative functions and social connections are redundant, but less when reproduction, leadership, ecological knowledge, or parental care are concentrated in particular individuals (Lukas & Clutton-Brock, 2012; Mech, 2000; Modlmeier et al., 2014). Ecological context may further modify these relationships because resource availability, environmental disturbance, population density, and opportunities for dispersal or breeder replacement can influence both group stability and the costs and benefits of remaining in social groups (Maldonado-Chaparro & Chaverri, 2021). Comparative studies across avian and mammalian cooperative breeders should therefore distinguish the number of potential helpers from their capacity to functionally replace the individuals lost.

## Acknowledgements

We thank the Oregon Department of Fish and Wildlife (ODFW) for long-term wolf monitoring and for supporting the collection of field data used in this study. We are especially grateful to the many current and former ODFW biologists and field staff who contributed to wolf monitoring and data collection during the study period. Additionally, we thank the field staff of the U.S. Fish and Wildlife Service, the Confederated Tribes of Warm Springs, and the Confederated Tribes of the Umatilla Indian Reservation for their contributions to wolf monitoring and data collection.

## Author Contributions

C. Aaron Bott conceived the ideas, designed the study, curated and analyzed data, and led the writing of the manuscript. Daniel R. MacNulty contributed to study design, interpretation of the analyses, and critically revised the manuscript. Roblyn Brown supervised long-term field data collection and critically reviewed the manuscript. All authors approved the final version for publication and agree to be accountable for the work.

## Statement of Inclusion

This study was conducted within Oregon, USA, and includes authors affiliated with institutions actively involved in wolf management within the study region. The research design and long-term monitoring program were developed in collaboration with state and federal wildlife agencies responsible for wolf conservation. Field data were collected through coordinated efforts among agency biologists, Tribal authorities, and local wildlife professionals. Relevant literature produced by researchers working within the region was cited where appropriate. Study findings will be shared with state and federal managers to inform ongoing conservation and management efforts.

## Data Availability

Data available from the Dryad Digital Repository: https://doi.org/10.5061/dryad.6m905qgq (Bott et al., 2026).

---

**Table S1.** Results from a logistic generalized linear mixed model assessing the effects of parental death, documented non-parental mortality, and pack size on the probability of wolf pack dissolution in Oregon, USA. Estimate refers to the log-odds coefficient; Std. Error is its standard error; z is the Wald test statistic; and p is the associated p-value. Odds ratios (OR) indicate how the odds of dissolution change with each predictor. Values above 1 indicate increased odds of dissolution, whereas values below 1 indicate decreased odds. The 95% confidence intervals (Lower and Upper 95% CI) show the likely range of values for each odds ratio.

| Predictor | Estimate | Std. Error | z | p | OR | Lower 95% CI | Upper 95% CI |
| --- | --- | --- | --- | --- | --- | --- | --- |
| (Intercept) | -0.92 | 0.54 | -1.70 | 0.089 | 0.40 | 0.14 | 1.15 |
| Parental death | 1.58 | 0.42 | 3.74 | <0.001 | 4.86 | 2.12 | 11.14 |
| Any non-parent death | 0.03 | 0.49 | 0.07 | 0.944 | 1.03 | 0.40 | 2.68 |
| Pack size | -0.24 | 0.09 | -2.55 | 0.011 | 0.79 | 0.66 | 0.95 |

**Table S2.** Results from a logistic generalized linear mixed model assessing the effects of parental mortality severity (no parental death, single-parent death, biparental death) and pack size on the probability of wolf pack dissolution in Oregon, USA. Estimate refers to the log-odds coefficient; Std. Error is its standard error; z is the Wald test statistic; and p is the associated p-value. Odds ratios (OR) indicate how the odds of dissolution change relative to packs in which both parents survived. The 95% confidence intervals (Lower and Upper 95% CI) show the likely range of values for each odds ratio.

| Predictor | Estimate | Std. Error | z | p | OR | Lower 95% CI | Upper 95% CI |
| --- | --- | --- | --- | --- | --- | --- | --- |
| (Intercept) | -0.79 | 0.54 | -1.45 | 0.148 | 0.46 | 0.16 | 1.32 |
| Single-parent death | 1.00 | 0.51 | 1.97 | 0.049 | 2.72 | 1.00 | 7.37 |
| Biparental death | 3.02 | 0.73 | 4.16 | <0.001 | 20.52 | 4.94 | 85.28 |
| Pack size | -0.26 | 0.10 | -2.73 | 0.006 | 0.77 | 0.64 | 0.93 |

**Table S3.** Results from a logistic generalized linear mixed model assessing the effects of parental death, documented non-parental death, and pack size on the probability of reproductive failure in wolf packs in Oregon, USA. Reproductive failure was defined as fewer than two pups surviving by the time of annual monitoring (January–mid-March). Estimate refers to the log-odds coefficient; Std. Error is its standard error; z is the Wald test statistic; and p is the associated p-value. Odds ratios (OR) indicate how the odds of reproductive failure change with each predictor: values above 1 reflect increased odds, values below 1 reflect reduced odds. The 95% confidence intervals (Lower–Upper 95% CI) show the likely range of values for each odds ratio.

| Predictor | Estimate | Std. Error | z | p | OR | Lower 95% CI | Upper 95% CI |
| --- | --- | --- | --- | --- | --- | --- | --- |
| (Intercept) | -0.44 | 0.53 | -0.83 | 0.406 | 0.64 | 0.23 | 1.82 |
| Parental death | 2.80 | 0.56 | 4.99 | <0.001 | 16.51 | 5.48 | 49.69 |
| Any non-parent death | 0.89 | 0.41 | 2.19 | 0.029 | 2.43 | 1.10 | 5.37 |
| Pack size | -0.18 | 0.08 | -2.32 | 0.020 | 0.83 | 0.71 | 0.97 |

**Table S4.**
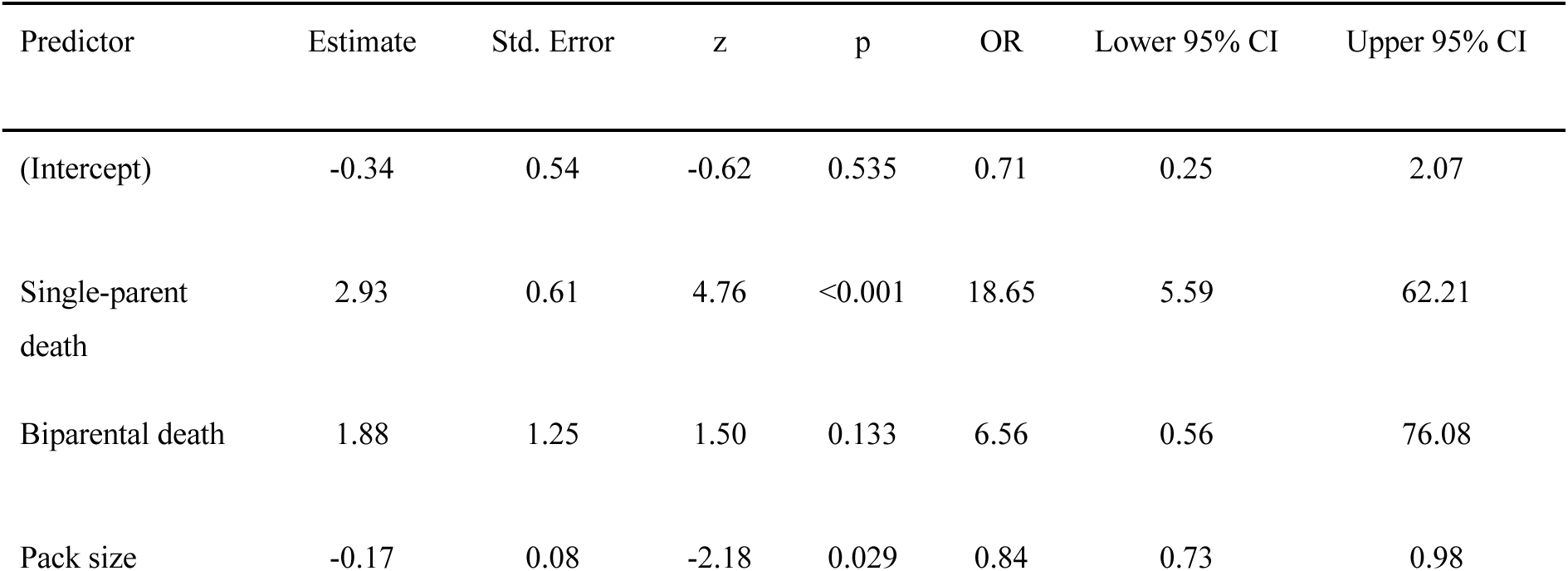
Results from a logistic generalized linear mixed model assessing the effects of single-parent death, biparental death, and pack size on the probability of reproductive failure in wolf packs in Oregon, USA. Reproductive failure was defined as fewer than two pups surviving by the time of annual monitoring (January–mid-March). Estimate refers to the log-odds coefficient; Std. Error is its standard error; z is the Wald test statistic; and p is the associated p-value. Odds ratios (OR) indicate how the odds of reproductive failure change with each predictor relative to packs in which both parents survived. The 95% confidence intervals (Lower–Upper 95% CI) show the likely range of values for each odds ratio.

| Predictor | Estimate | Std. Error | z | p | OR | Lower 95% CI | Upper 95% CI |
| --- | --- | --- | --- | --- | --- | --- | --- |
| (Intercept) | -0.34 | 0.54 | -0.62 | 0.535 | 0.71 | 0.25 | 2.07 |
| Single-parent death | 2.93 | 0.61 | 4.76 | <0.001 | 18.65 | 5.59 | 62.21 |
| Biparental death | 1.88 | 1.25 | 1.50 | 0.133 | 6.56 | 0.56 | 76.08 |
| Pack size | -0.17 | 0.08 | -2.18 | 0.029 | 0.84 | 0.73 | 0.98 |

**Table S5.** Results from a negative binomial generalized linear mixed model evaluating the effects of parental death, documented non-parental death, and pack size on pup recruitment in wolf packs in Oregon, USA. The response variable was the minimum number of pups surviving to 9–10 months. Estimate refers to the effect of each predictor on the log scale; Std. Error is the standard error of the estimate; z is the Wald test statistic; and p is the associated p-value. IRR (incidence rate ratio) indicates how the expected number of surviving pups changes with each predictor: values above 1 indicate more pups are expected, while values below 1 indicate fewer. The 95% confidence intervals (Lower and Upper 95% CI) show the likely range of values for each IRR.

| Predictor | Estimate | Std. Error | z | p | IRR | Lower 95% CI | Upper 95% CI |
| --- | --- | --- | --- | --- | --- | --- | --- |
| (Intercept) | 0.83 | 0.15 | 5.45 | <0.001 | 2.30 | 1.70 | 3.10 |
| Parental death | -1.09 | 0.21 | -5.10 | <0.001 | 0.34 | 0.22 | 0.51 |
| Any non-parent death | -0.19 | 0.12 | -1.62 | 0.104 | 0.83 | 0.66 | 1.04 |
| Pack size | 0.04 | 0.02 | 1.78 | 0.075 | 1.04 | 1.00 | 1.08 |

**Table S6.**
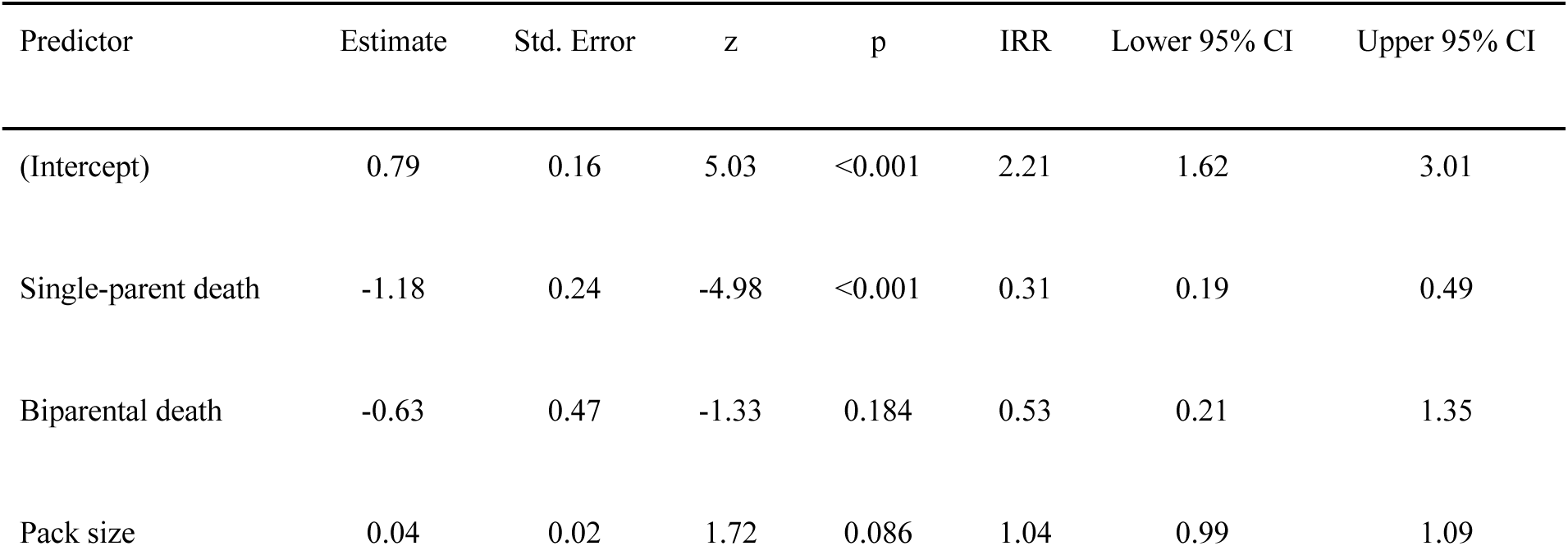
Results from a negative binomial generalized linear mixed model evaluating the effects of parental mortality severity (no parental death, single-parent death, biparental death) and pack size on pup recruitment in wolf packs in Oregon, USA. The response variable was the minimum number of pups surviving to 9–10 months. Estimate refers to the effect of each predictor on the log scale; Std. Error is the standard error of the estimate; z is the Wald test statistic; and p is the associated p-value. IRR (incidence rate ratio) indicates how the expected number of surviving pups changes relative to packs with no parental mortality. The 95% confidence intervals (Lower and Upper 95% CI) show the likely range of values for each IRR.

| Predictor | Estimate | Std. Error | z | p | IRR | Lower 95% CI | Upper 95% CI |
| --- | --- | --- | --- | --- | --- | --- | --- |
| (Intercept) | 0.79 | 0.16 | 5.03 | <0.001 | 2.21 | 1.62 | 3.01 |
| Single-parent death | -1.18 | 0.24 | -4.98 | <0.001 | 0.31 | 0.19 | 0.49 |
| Biparental death | -0.63 | 0.47 | -1.33 | 0.184 | 0.53 | 0.21 | 1.35 |
| Pack size | 0.04 | 0.02 | 1.72 | 0.086 | 1.04 | 0.99 | 1.09 |

